# VicMAG, an open-source tool for visualizing circular metagenome-assembled genomes highlighting bacterial virulence and antimicrobial resistance

**DOI:** 10.64898/2026.03.31.714378

**Authors:** Yusuke Tsuda, Yasuhiro Tanizawa, Thi My Hanh Vu, Yosuke Nishimura, Masaki Shintani, Haruka Abe, Futoshi Hasebe, Ikuro Kasuga, Miki Nagao, Masato Suzuki

**Affiliations:** Department of Clinical Laboratory Medicine, Kyoto University Graduate School of Medicine, 54 Shogoin-Kawahara-cho, Sakyo-ku, Kyoto, Kyoto 606-8507, Japan; Department of Infection Control and Prevention, Kyoto University Hospital, 54 Shogoin-Kawahara-cho, Sakyo-ku, Kyoto, Kyoto 606-8507, Japan; Department of Genetics, School of Life Science, The Graduate University for Advanced Studies (SOKENDAI), 1111 Yata, Mishima, Shizuoka 411-8540, Japan; Department of Informatics, National Institute of Genetics, 1111 Yata, Mishima, Shizuoka 411-8540, Japan; Institute of Tropical Medicine, Nagasaki University, Nagasaki, Nagasaki, 852-8523, Japan; Department of Urban Engineering, School of Engineering, The University of Tokyo, 7-3-1 Hongo, Bunkyo, Tokyo 113-8656, Japan; Institute for Extra-cutting-edge Science and Technology Avant-garde Research of Life (X-star), Japan Agency for Marine-Earth Science and Technology (JAMSTEC), 2-15 Natsushima-cho, Yokosuka, Kanagawa 237-0061, Japan; Department of Engineering, Graduate School of Integrated Science and Technology, Shizuoka University, 3-5-1 Johoku, Chuo-ku, Hamamatsu, Shizuoka 432-8561, Japan; Japan Collection of Microorganisms, RIKEN BioResource Research Center, 3-1-1, Koyadai, Tsukuba, Ibaraki 305-0074, Japan; Research Center for Advanced Science and Technology, The University of Tokyo, 4-6-1, Komaba, Meguro, Tokyo 153-8904, Japan; Antimicrobial Resistance Research Center, National Institute of Infectious Diseases, Japan Institute for Health Security, 4-2-1 Aobacho, Higashimurayama, Tokyo 189-0002, Japan

## Abstract

Bacterial pathogens spread in clinical and environmental settings, and mobile genetic elements (MGEs), such as plasmids and phages, mediate the transfer of virulence factor genes (VFGs) and antimicrobial resistance genes (ARGs) among bacterial communities. Metagenomic analysis of environmental and wastewater samples using highly accurate long-read sequencing technologies, such as PacBio HiFi sequencing, provides valuable insights into monitoring the regional spread of VFGs and ARGs, including dissemination mediated by MGEs. No visualization tool is currently available for the comprehensive display of numerous resulting circular metagenome-assembled genomes (cMAGs) with functional gene annotations. Here, we developed VicMAG, a visualization tool for highly complex cMAGs derived from long-read metagenome assemblies annotated using updated databases of VFGs, ARGs, and MGEs. Using 353 cMAGs from PacBio HiFi sequencing of a wastewater sample, we demonstrated the utility of VicMAG for metagenome visualization. VicMAG provides comprehensive, size-aware visualization of cMAGs representing bacterial chromosomes and plasmids, annotated with VFGs, ARGs, and phages. By simultaneously visualizing all cMAGs in a framework, VicMAG facilitates a holistic understanding of the distribution and genomic context of VFGs and ARGs across complex microbial communities. This tool supports integrated surveillance of bacteria associated with virulence and antimicrobial resistance across clinical, environmental, and One Health contexts.

## Introduction

Antimicrobial Resistance (AMR) is an urgent public health challenge globally. An estimated 8.22 million deaths were associated with bacterial AMR in 2050 [1]. Antimicrobial resistance genes (ARGs) spread among bacteria via mobile genetic elements (MGEs), such as plasmids, phages, insertion sequences, transposons, and integrons. Plasmids are major drivers of horizontal gene transfer of virulence factor genes (VFGs) and ARGs between bacteria. Continuous monitoring of this dissemination is crucial for public health to understand the mechanisms underlying AMR spread and to prepare for future public health crises.

Environmental and wastewater surveillance, often utilizing metagenomic analysis, is a resource-efficient approach for monitoring AMR [2–4]. Monitoring hospital wastewater is particularly important because untreated effluent contains high levels of bacteria harboring VFGs and ARGs, and its investigation can provide a regional snapshot of AMR [3].

Long-read sequencing technologies, such as Pacific Biosciences (PacBio) sequencing and Oxford Nanopore Technologies (ONT) nanopore sequencing, are effective for metagenomic analysis and significantly improve the quality of metagenome-assembled genomes (MAGs) [5]. PacBio HiFi sequencing is known for its high nucleotide accuracy, approximately 99.9% [6]. Unlike early-generation ONT reads, which exhibited relatively high error rates, PacBio HiFi reads achieve high accuracy while maintaining long read lengths [7].

Metagenome assemblers specialized for PacBio HiFi sequencing data include hifiasm-meta and metaMDBG [6, 8]. Among these, metaMDBG has been reported to be highly scalable and substantially faster than hifiasm-meta while requiring less memory [6]. Furthermore, metaMDBG demonstrated better recovery of high-quality circular MAGs (cMAGs) representing bacterial chromosomes and plasmids compared with hifiasm-meta [6]. MAGs recovered from metagenomic analysis often undergo binning and refinement for detailed characterization [9]. For visualizing these MAGs, graphical tools, such as Bandage and Circos, are available [10, 11]. Bandage is effective for visualizing numerous de novo assembly graphs across diverse genome assembly contexts, including metagenomic assemblies. However, it cannot display detailed genetic information such as VFGs and ARGs. Although Circos is useful for visualizing relationships between genomic intervals in a circular ideogram layout, it does not readily support comprehensive visualization of numerous annotated cMAGs.

Recently, tools such as GenoVi have emerged that use Circos to create custom circular genome representations of multiple genomes for comparative genomics studies [12]. While users can manually create scripts to generate such maps, tools like Circos often require programming skills, raising the barrier to entry for graphical visualization and making consistent comparisons difficult. Therefore, no existing tool readily provides effective and comprehensive visualization of genetic information derived from metagenomic analyses. In metagenomic datasets, hundreds of cMAGs representing chromosomes and plasmids may be recovered simultaneously. Because VFGs and ARGs can be unevenly distributed across diverse genetic backgrounds and genome size classes, examining only selected genomes may obscure important patterns of co-occurrence and size-dependent bias. Therefore, comprehensive visualization of all recovered cMAGs within a unified framework is essential for accurately interpreting the overall structure of VFGs, ARGs, and MGEs across complex microbial communities.

In this study, we developed a tool called Visualization of circular Metagenome-Assembled Genome (VicMAG) to comprehensively visualize cMAGs representing bacterial chromosomes and plasmids annotated with VFGs, ARGs, and phage genes derived from metagenomic assemblies. We demonstrated visualization by VicMAG of 353 cMAGs assembled from PacBio HiFi sequencing of a wastewater sample. As a unique feature, VicMAG enables the simultaneous visualization of bacterial and plasmid genomes with functional annotations based on updated databases of these genetic elements, thereby supporting comprehensive analysis of metagenomic datasets.

## Materials and Methods

### Working flow

VicMAG requires cMAGs generated by the long-read assemblers hifiasm-meta or metaMDBG as input prior to visualization (Figure 1A). hifiasm-meta and metaMDBG indicate putative circularity in their output files: hifiasm-meta appends the suffix “c” to contig names, whereas metaMDBG appends “circular = yes” to MAG descriptions. After assembly, cMAGs marked with the suffix “c” or ʻcircular = yes’ are manually extracted, and coding sequences (CDSs) are predicted and annotated using DFAST [13] with the VFG and ARG detection option enabled, based on the Virulence Factor Database (VFDB) and the Comprehensive Antibiotic Resistance Database (CARD), respectively [14, 15]. Optionally, cMAGs are classified as plasmids using geNomad [16], which simultaneously detects phage regions. Following data preprocessing, VicMAG visualizes the cMAGs. In this study, hifiasm-meta v0.13, metaMDBG v1.0, DFAST v1.3.4, CARD v3.2.9, VFDB v2024-05-10, and geNomad v1.11.1 were used. Thresholds for identification of VFGs and ARGs were used in the default settings: ≥90% identity and ≥90% coverage. The threshold for identification of phage genomes and plasmids was used in the default settings: a score of ≥0.70.

**Figure 1.**
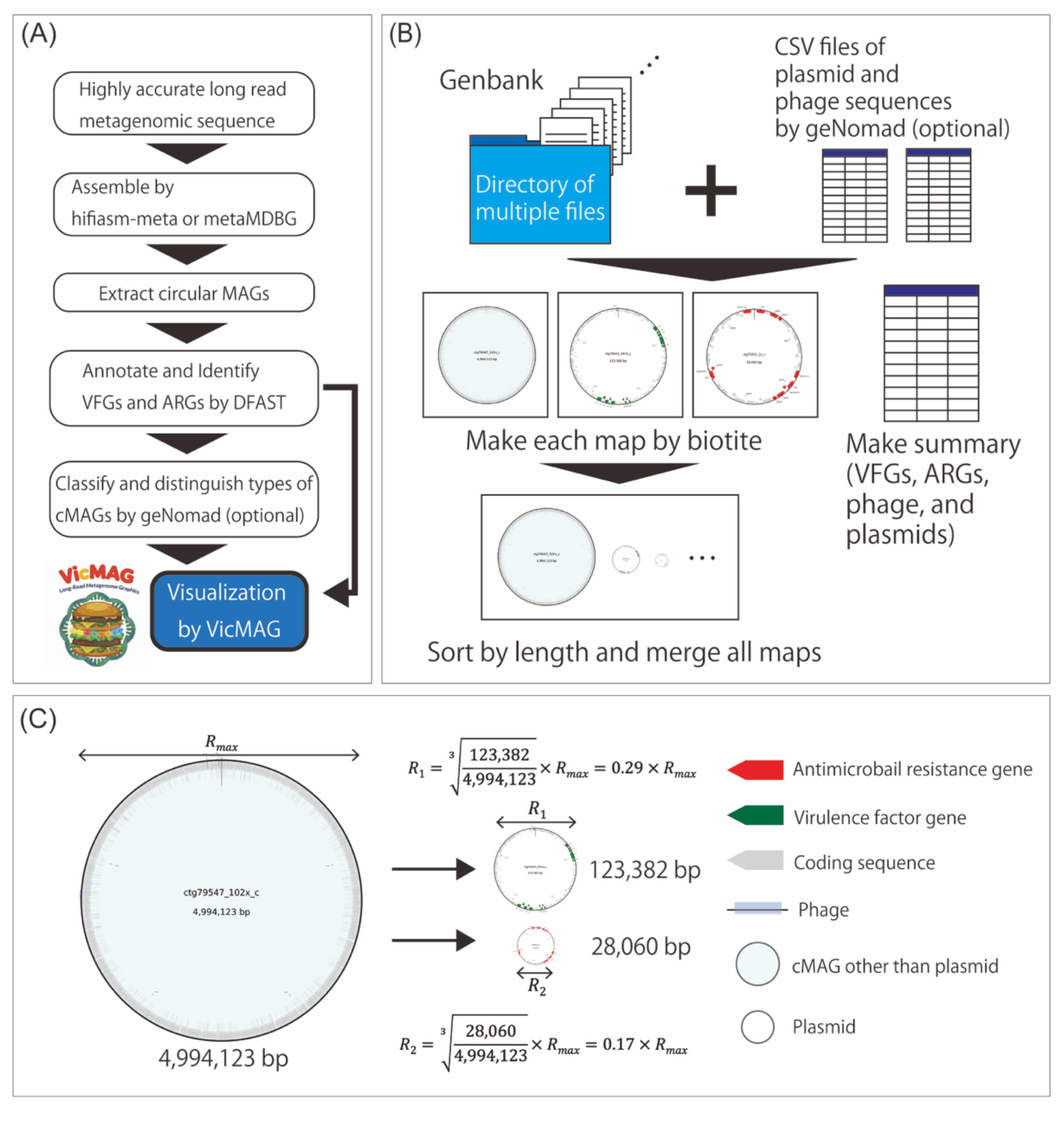
Overview of VicMAG and processing. (A) To visualize data with VicMAG, several steps are required. Highly accurate long reads are first assembled using hifiasm-meta or metaMDBG to obtain MAGs, from which cMAGs are extracted. These cMAGs are annotated with DFAST, including identification of VFGs and ARGs. Optionally, geNomad is used to classify cMAGs as plasmid or non-plasmid. The processed data are then visualized using VicMAG. (B) VicMAG requires a directory containing multiple GenBank files and optional CSV files containing plasmid and phage information. When their file paths are provided via the command line, VicMAG generates and stores individual maps for cMAGs in an output directory, along with a summary file. The maps are subsequently sorted by sequence length and merged into a unified visualization. (C) Because cMAGs vary greatly in size, VicMAG applies cubic ratio scaling to normalize their diameters for display. The largest cMAG is assigned a maximum radius (*R_max_*), and the others are scaled accordingly. Each map shows VFGs, ARGs, coding sequences, and phage genomes. Plasmid cMAGs are displayed without color, whereas non-plasmid cMAGs are filled with cyan for clear differentiation.

VicMAG is implemented in Python 3. VicMAG generates graphical maps from multiple GenBank files of cMAGs stored in a working directory (Figure 1B). For each nucleotide sequence (e.g., chromosome or plasmid), it constructs a map displaying DFAST-annotated VFGs and ARGs, together with the geNomad-detected genomic positions of phage sequences and plasmid classifications, using the Biotite library (https://github.com/biotite-dev/biotite). VicMAG generates maps for each cMAG individually and then combines them into a single overview map after adjusting margins and layout. Each individual cMAG map is also saved in the output directory, allowing users to inspect individual cMAGs separately when detailed examination is required. To improve readability, label overlap is reduced by shifting individual labels. In addition, VicMAG exports a CSV file summarizing the identified VFGs, ARGs, phage sequences, and plasmid classifications. The number of cMAGs displayed per column can be freely adjusted by the user via command-line options. Users can choose to display exclusively plasmid-derived cMAGs containing VFGs and/or ARGs, with this filter also being fully customizable through command-line options.

VicMAG is available from the GitHub repository (https://github.com/TsudaYusuke/VicMAG) and can be used either by directly downloading the required files from GitHub or by installing the package via Bioconda. Detailed installation procedures, example commands, and usage instructions are provided in the GitHub repository.

### Design

cMAGs consist of chromosomes, plasmids, and other elements, and their lengths differ by over 1,000-fold. To visualize all cMAGs in a single figure, a cubic scaling approach was applied. Among all cMAGs, the one with the longest sequence length (*L_max_*) was used as the reference, and its diameter in the map was defined as *R_max_*. If *L_each_* and *R_each_* represent the sequence length and corresponding diameter in the map, respectively, *R_each_* was calculated as:

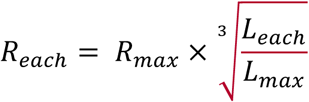

After calculating *R_each_*, all cMAGs are mapped in a single figure (Figure 1C). VFGs and ARGs are colored red and green, respectively. The gene names of VFGs and ARGs are displayed, whereas those of CDSs are not shown. In chromosomes, regions containing more than 8 VFGs within 5,000 bp are clustered and displayed as arcs outside the map together with a list of VFGs. Phage sequences are colored light purple. cMAGs not classified as plasmids by geNomad are filled with light cyan. All colors are configurable via command-line options.

### Sample

To demonstrate the mapping of cMAGs using VicMAG, a water sample collected from a polluted urban river near hospitals in Hanoi, Vietnam, in 2021 was analyzed. Bacteria were cultured in Tryptic Soy Broth (BD Difco, USA) containing 4 mg/L colistin (Sigma-Aldrich, USA), a last-resort antibiotic used to enrich colistin-resistant bacteria. High-molecular-weight DNA was extracted using an enzymatic method. Metagenomic HiFi sequencing was performed on the Sequel II system (PacBio, USA). The raw sequencing reads were deposited in the NCBI Sequence Read Archive (SRA) under accession number DRX553298.

Metagenomic HiFi sequencing reads were assembled using hifiasm-meta or metaMDBG with default parameters. The resulting cMAGs were subsequently annotated using DFAST for the identification of VFGs and ARGs. In addition, geNomad was employed with default parameters to classify the cMAGs as plasmids or non-plasmids and to detect phage sequences.

## Results

### Summary of metagenome assembly

A total of 40.4 Gb of metagenomic HiFi sequencing data was obtained from the colistin-enriched culture. Assembly using hifiasm-meta and metaMDBG yielded 44,300 and 28,857 MAGs, respectively (Supplementary Table 1; see fasta files available on Figshare). Taxonomic classification of the 28,857 metaMDBG-derived MAGs using Kraken2 v2.0.8 revealed diverse microbial taxa, including Enterobacterales, which frequently spread VFGs and ARGs via MGEs (Supplementary Figure 1) [17]. The numbers of cMAGs were 347 and 353 for hifiasm-meta and metaMDBG (see fasta files available on Figshare), respectively. The cMAGs generated by hifiasm-meta and metaMDBG were compared based on the coverage profile of each cMAG. As a result, 230 cMAGs, representing 65.2% of the cMAGs generated by metaMDBG, showed high similarity to those generated by hifiasm-meta (Supplementary Figure 2 and Supplementary Data). For subsequent analyses, the 353 metaMDBG-derived cMAGs were used (Figure 2). The median length of the cMAGs was 52,962.0 bp (interquartile range 41,184.5‒74,635.5 bp), with a minimum of 10,910 bp and a maximum of 4,079,750 bp. Among these, 93 cMAGs harbored known ARGs, including six mobile colistin resistance (*mcr*) genes, and 25 harbored known VFGs, including *pilA* and *iro* genes (Supplementary Data). A total of 226 cMAGs were classified as plasmids (size range: 10,910‒280,267 bp), representing diverse plasmid types, including *mcr*-carrying MOBP1 plasmids, whereas 11 cMAGs, classified as 10 chromosomes and one plasmid, contained phage sequences (Supplementary Data).

**Figure 2.**
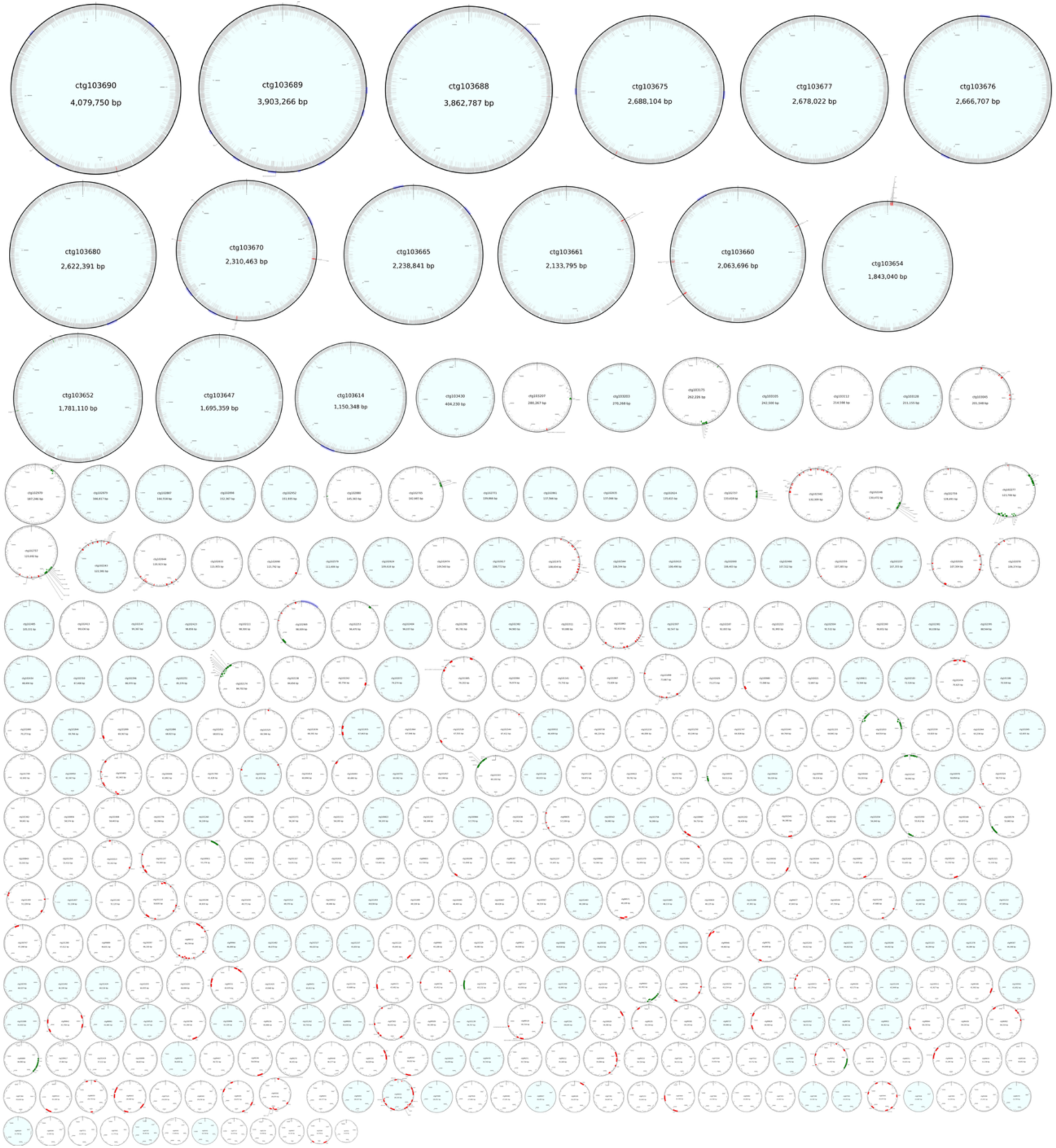
Visualization by VicMAG and maps of 353 cMAGs assembled by metaMDBG. cMAGs obtained from hospital effluent cultures grown in the presence of colistin were visualized using VicMAG.

### Visualization of cMAGs using VicMAG

As designed, all 353 metaMDBG-derived cMAGs, including 226 plasmids, were visualized using VicMAG (Figure 2; see the high-resolution figure available on Figshare), highlighting cMAGs containing VFGs (red), ARGs (green), and/or phage sequences (light purple). Non-plasmid cMAGs were filled with light cyan. The cMAGs were sorted in descending order of sequence length and arranged such that the six largest cMAGs were placed in the top row, resulting in a total of 18 rows. The wall-clock runtime was 154.9 seconds, and the peak memory usage was approximately 8.4 GB. Among the generated cMAGs, three derived from gram-negative bacteria, as classified by Kraken2, were selected for detailed visualization in Figure 3 [17]. ctg103689, ctg103045, and ctg101474 were assigned to *Morganella morganii*, *Escherichia coli*, and *E. coli*, respectively. geNomad classified ctg103045 and ctg101474 as plasmids, and identified seven phage genomes in ctg103689. Additionally, 226 cMAGs were identified as plasmids by geNomad and visualized using VicMAG (Supplementary Figure 3; see the high-resolution figure available on Figshare). By selectively displaying only representative or characteristic sequences, VicMAG provided a clearer visualization, enabling users to focus on the meaningful features while maintaining clarity and interpretability in large-scale metagenomic datasets.

**Figure 3.**
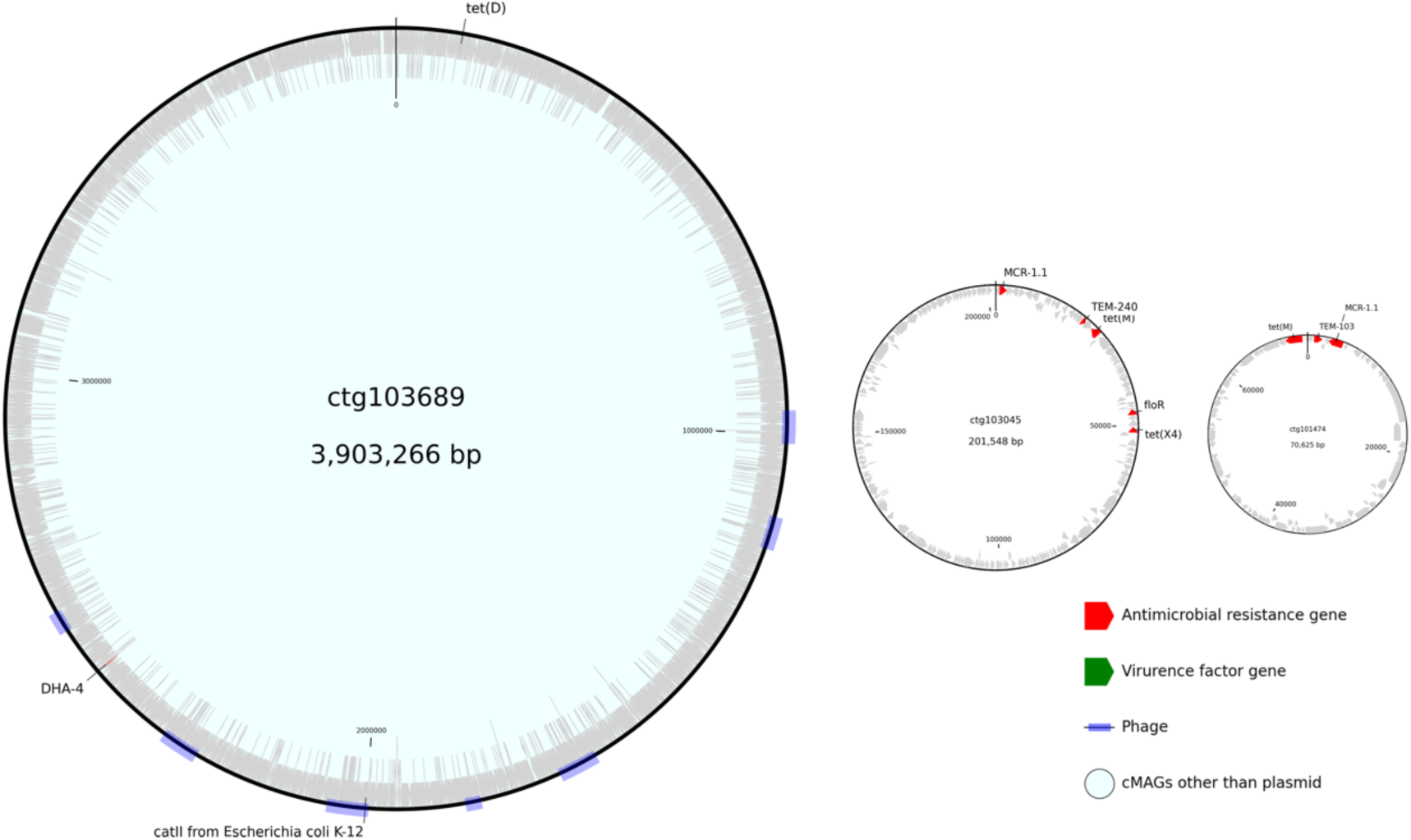
Visualization of three cMAGs derived from gram-negative bacteria carrying phage genomes and antimicrobial resistance genes. Three cMAGs carrying phage genomes and antimicrobial resistance genes including *mcr* genes, which confer resistance to colistin, were mapped by VicMAG.

## Discussion

We developed VicMAG, a visualization tool that enables comprehensive mapping of numerous cMAGs derived from metagenomic analyses, with a particular focus on VFGs and ARGs, and demonstrated its utility in large-scale metagenomic datasets. Existing visualization tools are generally limited to comparing a small number of similar sequences or displaying genomes of varying sizes [11, 12, 18]. However, cMAGs derived from metagenomic datasets often exhibit substantial diversity in their genetic composition, which limits the effectiveness of such tools for large-scale comparative visualization. In large-scale metagenomic studies, interpreting the ecological and epidemiological significance of VFGs and ARGs requires identifying their presence and understanding their distribution across all genetic elements within microbial communities. Simultaneous visualization of all cMAGs allows researchers to distinguish widespread dissemination from rare or limited distribution and to evaluate whether VFGs, ARGs, and MGEs are predominantly associated with plasmids, chromosomes, or specific genome size ranges. Moreover, as cMAG sizes can vary by up to three orders of magnitude, appropriate size normalization is essential for an informative overview. While GenoVi employs a square-root scaling approach for genome size normalization [12], this approach was insufficient for metagenomic analyses, in which genome sizes differed by approximately three orders of magnitude. Therefore, VicMAG adopts a cubic scaling method to provide a more balanced and interpretable visual representation.

In contrast to several existing tools that incorporate additional information such as GC skew or tRNA features [11, 12], VicMAG omits these elements to reduce visual complexity and maintain focus on VFGs, ARGs, plasmids, and phages. In the current implementation, analysis results are provided as static images. VicMAG visualizes cMAGs based on the outputs of assembly and annotation tools, and does not independently validate circularity or assess potential misassemblies. Therefore, the interpretation of gene co-localization and genomic context should be made with consideration of the quality and limitations of the underlying assembly outputs. Overall, the current release focuses on essential functionalities, providing a simplified framework for intuitive visualization.

VicMAG utilizes analysis results from DFAST and geNomad, and continued improvements in functional annotations by these tools are expected to further strengthen VicMAG’s visualization capabilities. In addition to VFG and ARG annotation, DFAST is expected to further enhance its annotation of MGEs, such as insertion sequences, transposons, and integrons, as well as genes involved in defense and anti-defense systems against phages and plasmids. Once these functionalities are implemented, VicMAG will be updated to support visualization of the resulting annotations. In addition, species identification of non-plasmid cMAGs using DFAST_QC [19] would enable species-level information to be displayed on cMAGs in VicMAG visualizations. Furthermore, integrating analyses currently performed outside VicMAG into its internal pipeline may improve usability and streamline the user workflow.

The present study was mainly based on a single colistin-enriched wastewater dataset. We also analyzed a meropenem-enriched dataset generated from the same wastewater sample (accession number DRR569829), and the additional results have been deposited on Figshare. However, the use of antimicrobial-enriched datasets may introduce selection bias and may not fully represent typical microbial communities across different environments. Therefore, the observed distributions of VFGs, ARGs, and MGEs should be interpreted with caution when considering their generalizability.

In this study, assemblies generated from PacBio HiFi sequencing reads were used for analysis. VicMAG can also visualize assemblies derived from ONT sequencing reads, provided that they are generated using metaMDBG (v1.0 or later). Furthermore, VicMAG enables the integrated visualization of multiple GenBank files, highlighting genomic features such as VFGs, ARGs, plasmids, and phage-related sequences. In the future, we aim to extend VicMAG to support complete genome assemblies of individual strains derived from metagenomic binning or single-cell long-read sequencing. This versatility underscores the broad applicability of VicMAG across diverse genomic and metagenomic datasets.

## Data Availability

The source code for VicMAG is publicly available on GitHub (https://github.com/TsudaYusuke/VicMAG) and Figshare (https://doi.org/10.6084/m9.figshare.31841554). MAGs, their summaries, and high-resolution images are publicly available on Figshare (https://doi.org/10.6084/m9.figshare.31841554).

## Acknowledgements

We thank the Information Technology Center at Nagoya University, Japan, for providing computational resources on the supercomputer "Flow".

## Supplementary Materials

Supplementary Table 1.

Supplementary Figures 1, 2, and 3.

Supplementary Data.

## References

1. Collaborators GBDAR: Global burden of bacterial antimicrobial resistance 1990-2021: a systematic analysis with forecasts to 2050. Lancet 2024, 404(10459):1199-1226.

2. Larsson DGJ, Flach CF, Laxminarayan R: Sewage surveillance of antibiotic resistance holds both opportunities and challenges. Nat Rev Microbiol 2023, 21(4):213–214.

3. Philo SE DLK, Noble RT, Zhou NA, Alghafri R, Bar-Or I, Darling A, D’Souza N, Hachimi O, Kaya D, Kim S, Gaardbo Kuhn K, Layton BA, Mansfeldt C, Oceguera B, Radniecki TS, Ram JL, Saunders LP, Shrestha A, Stadler LB, Steele JA, Stevenson BS, Vogel JR, Bibby K, Boehm AB, Halden RU, Delgado Vela J.: Wastewater surveillance for bacterial targets current challenges and future goals. Appl Environ Microbiol 2024, 90(1).

4. Berglund F, Ebmeyer S, Kristiansson E, Larsson DGJ: Evidence for wastewaters as environments where mobile antibiotic resistance genes emerge. Commun Biol 2023, 6(1):321.

5. Espinosa E, Bautista R, Larrosa R, Plata O: Advancements in long-read genome sequencing technologies and algorithms. Genomics 2024, 116(3):110842.

6. Benoit G, Raguideau S, James R, Phillippy AM, Chikhi R, Quince C: High-quality metagenome assembly from long accurate reads with metaMDBG. Nat Biotechnol 2024, 42(9):1378–1383.

7. Kim CY, Ma J, Lee I: HiFi metagenomic sequencing enables assembly of accurate and complete genomes from human gut microbiota. Nat Commun 2022, 13(1):6367.

8. Feng X, Cheng H, Portik D, Li H: Metagenome assembly of high-fidelity long reads with hifiasm-meta. Nat Methods 2022, 19(6):671–674.

9. Mallawaarachchi V, Wickramarachchi A, Xue H, Papudeshi B, Grigson SR, Bouras G, Prahl RE, Kaphle A, Verich A, Talamantes-Becerra B et al: Solving genomic puzzles: computational methods for metagenomic binning. Brief Bioinform 2024, 25(5).

10. Wick RR, Schultz MB, Zobel J, Holt KE: Bandage: interactive visualization of de novo genome assemblies. Bioinformatics 2015, 31(20):3350–3352.

11. Krzywinski M, Schein J, Birol I, Connors J, Gascoyne R, Horsman D, Jones SJ, Marra MA: Circos: an information aesthetic for comparative genomics. Genome Res 2009, 19(9):1639–1645.

12. Cumsille A, Duran RE, Rodriguez-Delherbe A, Saona-Urmeneta V, Camara B, Seeger M, Araya M, Jara N, Buil-Aranda C: GenoVi, an open-source automated circular genome visualizer for bacteria and archaea. PLoS Comput Biol 2023, 19(4):e1010998.

13. Tanizawa Y, Fujisawa T, Nakamura Y: DFAST: a flexible prokaryotic genome annotation pipeline for faster genome publication. Bioinformatics 2018, 34(6):1037–1039.

14. Alcock BP, Huynh W, Chalil R, Smith KW, Raphenya AR, Wlodarski MA, Edalatmand A, Petkau A, Syed SA, Tsang KK et al: CARD 2023: expanded curation, support for machine learning, and resistome prediction at the Comprehensive Antibiotic Resistance Database. Nucleic Acids Res 2023, 51(D1):D690-D699.

15. Zhou S, Liu B, Zheng D, Chen L, Yang J: VFDB 2025: an integrated resource for exploring anti-virulence compounds. Nucleic Acids Res 2025, 53(D1):D871-D877.

16. Camargo AP, Roux S, Schulz F, Babinski M, Xu Y, Hu B, Chain PSG, Nayfach S, Kyrpides NC: Identification of mobile genetic elements with geNomad. Nat Biotechnol 2024, 42(8):1303–1312.

17. Wood DE, Lu J, Langmead B: Improved metagenomic analysis with Kraken 2. Genome Biol 2019, 20(1):257.

18. Grant JR, Enns E, Marinier E, Mandal A, Herman EK, Chen CY, Graham M, Van Domselaar G, Stothard P: Proksee: in-depth characterization and visualization of bacterial genomes. Nucleic Acids Res 2023, 51(W1):W484–W492.

19. Elmanzalawi M, Fujisawa T, Mori H, Nakamura Y, Tanizawa Y: DFAST_QC: quality assessment and taxonomic identification tool for prokaryotic Genomes. BMC Bioinformatics 2025, 26(1):3.

