## Supplementary Materials for "VicMAG, an open-source tool for visualizing circular metagenome-assembled genomes highlighting bacterial virulence and antimicrobial resistance"

**Supplementary Table 1. Comparison of assembly statistics generated by hifiasm-meta and metaMDBG using PacBio HiFi sequencing data (SRA accession: DRX553298)**

| | hifiasm-meta | metaMDBG |
|---|---|---|
| Assembled metagenome size (Gb) | 2.16 | 1.28 |
| No. of contigs (MAGs) | 44,300 | 28,857 |
| No. of circular contigs (cMAGs) | 347 | 353 |
| No. of near-complete MAGs* | 29 | 20 |
| No. of >1Mb near-complete cMAGs* | 23 | 13 |
| No. of cMAGs classified as plasmids† | 202 | 226 |
| Peak memory usage (GB) | 204.56 | 10.4 |
| Wall-clock time (h) | 23 | 5 |

*Assessed by CheckM v1.1.0 [1]. Near-complete: ≥90% completeness and ≤5% contamination

† Assessed by geNomad v1.11.1 [2]

Abbreviation: MAG, metagenome-assembled genome; cMAG, circular metagenome-assembled genome

**Supplementary Figure 1. Result of Kraken2 analysis of MAGs assembled by metaMDBG from PacBio HiFi sequencing data (SRA accession: DRX553298)**

PacBio HiFi sequencing data obtained from wastewater cultures with colistin were assembled using metaMDBG. All resulting MAGs were taxonomically analyzed with Kraken2 v2.1.1 (database: standard 2025) [3]. Taxonomic abundance was subsequently estimated using Bracken v3.0.1, and the taxonomic composition was visualized with Krona v2.8.1 [4, 5]. This figure is available on Figshare (https://doi.org/10.6084/m9.figshare.31841554).

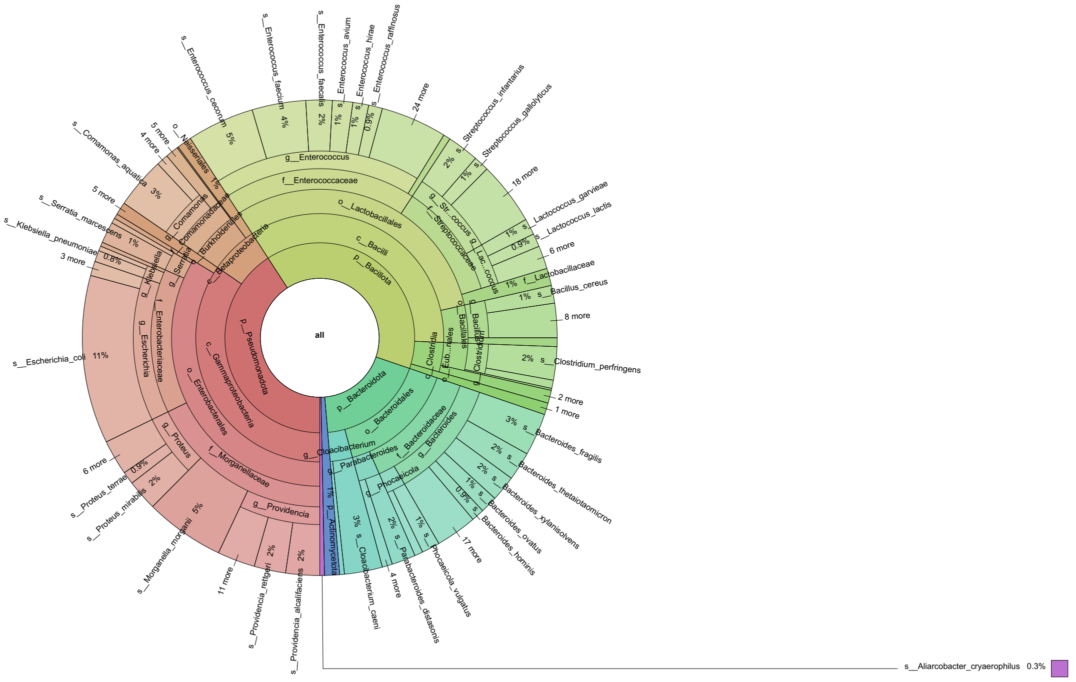

**Supplementary Figure 2. Relation of cMAGs generated by hifiasm-meta and metaMDBG**

Using hifiasm-meta, 347 cMAGs were generated, whereas metaMDBG generated 353 cMAGs. These cMAGs were analyzed using comprehensive BLASTN searches to identify homologous regions among the sequences. The coverage of each cMAG was calculated based on these homologous regions, and connections between cMAGs were visualized in a Sankey chart for pairwise comparisons with >80% coverage (see the HTML file of the Sankey chart available on Figshare). The cMAGs from metaMDBG were sorted by sequence length. Overall, 250 cMAGs (72.0%) from hifiasm-meta and 230 cMAGs (65.2%) from metaMDBG were connected.

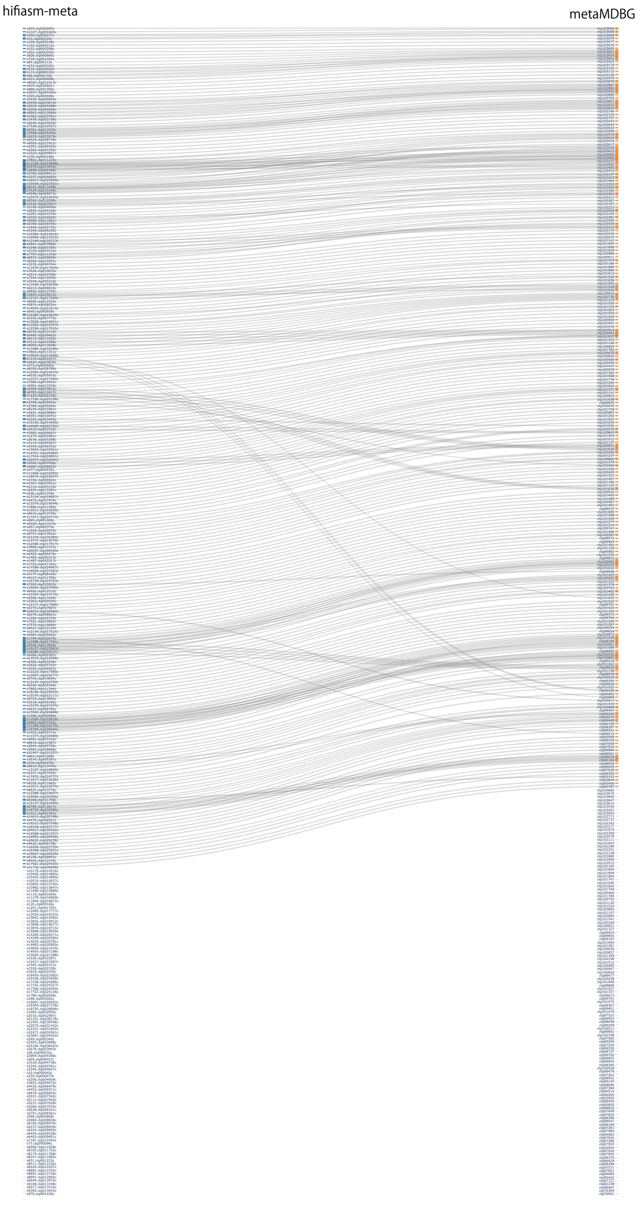

**Supplementary Figure 3. Visualization of cMAGs classified as plasmids using VicMAG**

Using the optional command, 226 cMAGs classified as plasmids were selectively visualized with VicMAG.

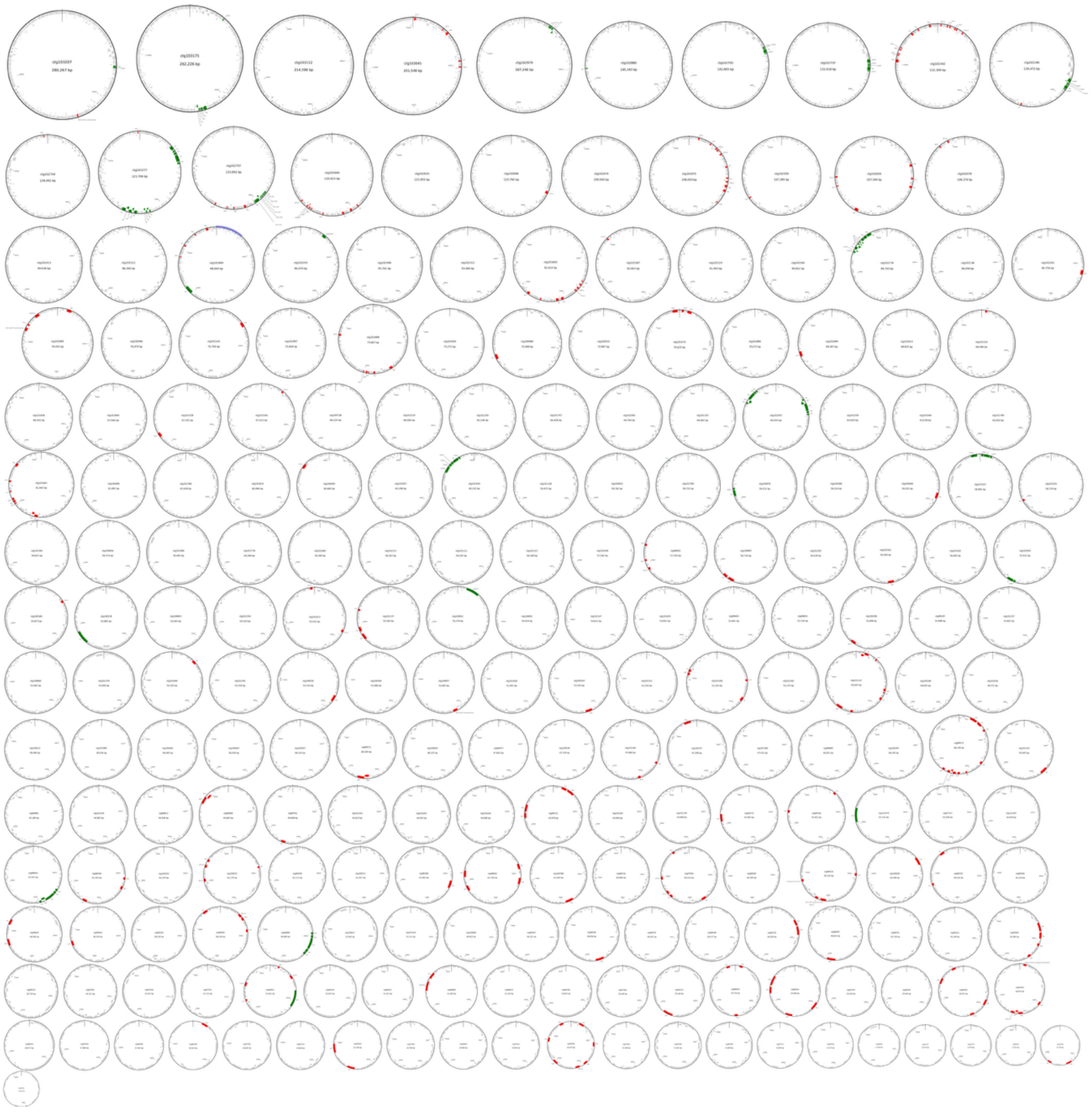

### References


1. Parks DH, Imelfort M, Skennerton CT, Hugenholtz P, Tyson GW: **CheckM: assessing the quality of microbial genomes recovered from isolates, single cells, and metagenomes**. *Genome Res* 2015, **25**(7):1043-1055.
2. Camargo AP, Roux S, Schulz F, Babinski M, Xu Y, Hu B, Chain PSG, Nayfach S, Kyrpides NC: **Identification of mobile genetic elements with geNomad**. *Nat Biotechnol* 2024, **42**(8):1303-1312.
3. Wood DE, Lu J, Langmead B: **Improved metagenomic analysis with Kraken 2**. *Genome Biol* 2019, **20**(1):257.
4. Lu J, Breitwieser FP, Thielen P, Salzberg SL: **Bracken: estimating species abundance in metagenomics data**. *PeerJ Comput Sci* 2017, **3**.
5. Ondov BD BN, Phillippy AM. : **Interactive metagenomic visualization in a Web browser**. *BMC Bioinformatics* 2011, **12:385.**
